# Effects of carbon nanotubes and derivatives of graphene oxide on soil bacterial diversity

**DOI:** 10.1101/591842

**Authors:** Christian Forstner, Thomas G. Orton, Peng Wang, Peter M. Kopittke, Paul G. Dennis

## Abstract

Carbon nanotubes (CNTs), reduced graphene oxide (rGO) and ammonia-functionalized graphene oxide (aGO), are nanomaterials that possess varied and useful properties. However, following their use, their release into the environment is inevitable. While CNTs have been shown to influence soil bacterial diversity, albeit at very high concentration, the effects of rGO have only been examined using pure bacterial cultures, and those of aGO are unknown. Here, we investigated the effects of CNTs, rGO and aGO, at three time points (7, 14 and 30 days), and over a range of concentrations (1 ng, 1 µg and 1 mg kg dry soil^-1^), on soil bacterial diversity using 16S rRNA amplicon sequencing. Graphite was included to facilitate comparisons with a similar and naturally occurring carbon material, while the inclusion of GO allowed the effects of GO modification to be isolated. Bacterial community composition, but not alpha diversity, was altered by all treatments except the low GO, low rGO and high aGO treatments on day 14 only. In all cases, the nanomaterials led to shifts in community composition that were of similar magnitude to those induced by graphite and GO, albeit with differences in the taxa affected. Our study highlights that nanocarbon materials can induce changes in soil bacterial diversity, even at doses that are environmentally realistic.

## Introduction

Engineered carbon nanomaterials, such as carbon nanotubes (CNTs) and various forms of graphene oxide (GO), possess unique physical properties that facilitate their use in a wide-range of applications (Gao, 2015; Perreault et al., 2015; Ramakrishnan and Shanmugam, 2016; Taha and Alsharef, 2017; Xin et al., 2012). Models estimate that for CNTs, 0.004-1.6 μg kg^−1^ enter soils annually (Sun et al., 2014), with release rates of other carbon nanomaterials (e.g. fullerenes and GO) expected to be similar (Forstner et al., 2019; Sun et al., 2014). Recently, we have shown that relevant concentrations of GO (1 ng, 1 µg and 1 mg kg dry soil^-1^) significantly influence soil microbial community composition. However, these effects were of similar magnitude to those induced by equivalent doses of graphite – an analogous non-nanomaterial that occurs naturally in many soils (Forstner et al., 2019). At present, the effects of CNTs on soil bacterial diversity have only been investigated at concentrations that greatly exceed the estimated rates of release (6,250-2,500,000,000 times higher) (Chung et al., 2011; Ge et al., 2016; Jin et al., 2013, 2014; Shrestha et al., 2013; Sun et al., 2014), while those associated with GO derivatives are unknown. Given the important roles of bacteria in mediating many soil ecosystem services (Bardgett and van der Putten, 2014), the potential impacts of relevant doses of CNTs and GO derivatives on soil bacterial communities need to be examined. This information will help to facilitate safe and sustainable management of nanomaterials as their portfolio of applications continues to expand.

At extremely high concentration (≥30 mg kg^−1^ soil), the addition of CNTs to soil has been observed to reduce microbial biomass (Chung et al., 2011; Jin et al., 2013, 2014) and enzyme activity (Chung et al., 2011; Jin et al., 2013), and influence the structure of bacterial communities (Ge et al., 2016; Jin et al., 2014; Shrestha et al., 2013). These effects, however, have been shown to be similar to those induced by natural and industrial carbonaceous materials such as biochar and carbon black (Ge et al., 2016). Furthermore, at the lowest CNT concentration examined to date (10 mg kg^−1^ soil, *viz*. 6,250 times higher than the estimated rate of release), soil respiration, enzyme activity and bacterial community composition were not significantly affected (Shrestha et al., 2013). For derivatives of GO, such as reduced GO (rGO) and ammonia-functionalized GO (aGO), we are not aware of any previous studies that have examined their effects on soil microbial diversity, biomass or activity. However, in pure culture experiments, rGO has been shown to exhibit bactericidal properties (Guo et al., 2017; Gurunathan et al., 2012; Liu et al., 2011). No information is available regarding the interactions between bacteria and aGO.

In this study, we investigated the effects of CNTs, rGO and aGO on the diversity of soil bacterial communities at a range of environmentally relevant low, high and very high doses (1 ng, 1 µg and 1 mg kg dry soil^-1^) based on models of CNT release (Sun et al., 2014). To contextualize our findings, we included graphite-amended and GO-amended soils, as well as no-treatment controls (Forstner et al., 2019). The inclusion of graphite allowed us to interpret the effects of CNTs, rGO and aGO relative to a similar and naturally occurring carbonaceous material, while the inclusion of GO allowed any effects of GO modification to be isolated. Treatments were run in triplicate and bacterial communities were characterized after 7, 14 and 30 days using high-throughput 16S rRNA gene amplicon sequencing.

## 2. Materials and methods

### 2.1 Experimental design

A Kandosol (Australian Soil Classification; Isbell, 2002), otherwise known as an Ultisol (USDA Soil Taxonomy; Soil Survey Staff, 2014), was collected from 0-20 cm depth at a pineapple (*Ananas comosus*) farm in Queensland, Australia (27.02 °S, 152.92 °E). physicochemical properties of this soil have been described previously (Wang et al., 2016). Briefly, the soil was a sandy loam, with a pH of 5.4 (1:5, soil:water), an electrical conductivity of 1.0 dS/m (saturation extract), a cation exchange capacity of 2.6 cmol_c_/kg, and an organic C content of 1.1 % (Wang et al., 2016). Fresh soil was passed through a 2 mm sieve and then split into sub-samples to which the treatments were applied. Multi-walled carbon nanotubes (CNTs) (Sigma Aldrich, Cat: 698849-1G, Lot# MKBH5811), reduced graphene oxide (rGO) (Aldrich Chemistry, Cat: 777684-250MG, Lot# MKBR1638V), and ammonia functionalized graphene oxide (aGO) (Aldrich Chemistry, Cat: 791520-25ML, Lot# MKBR2522V) were applied to the soil at rates of 1 ng, 1 µg, and 1 mg kg dry soil^-1^ using a fine mist sprayer and thoroughly homogenized by mechanical mixing. All particles were dispersed into deionized water at a volume that resulted in the soil being adjusted to 50% water holding capacity (WHC). Three replicate 500 g samples of each treatment were placed into 1 L plastic containers with lids that facilitated gas exchange. This yielded 27 containers that were incubated for 30 days in the dark at 25°C, with the humidity maintained at 80% in order to keep the soils at the same WHC throughout the experimental period.

In the same experiment, we also established 1 ng, 1 µg, and 1 mg kg dry soil^−1^ graphite and graphene oxide (GO) treatments, as well as no-treatment controls, which were sprayed with an equal quantity of deionized water only and then mixed in the same way as all other treatments. Data for the no-treatment controls, and GO and graphite treatments have already been published (Forstner et al., 2019), but are included in our present study to contextualize our findings for the multiple doses of CNTs, rGO, and aGO.

### 2.2 Soil sampling and DNA extraction

Using sterile 50 ml plastic tubes, we collected c. 25 g soil cores from each experimental unit after 7, 14 and 30 days, and then transferred the soil to −80°C storage. All samples were then thawed and DNA was extracted from 250 mg soil using the Power Soil DNA Isolation kit (MO BIO Laboratories, Carlsbad, CA) according to the manufacturers’ instructions.

### 2.3 Bacterial 16S rRNA gene amplification, sequencing and analysis

Polymerase chain reaction (PCR) and sequencing 16S rRNA genes, and the subsequent bioinformatic analyses were performed as described in our previous study (Forstner et al., 2019). Briefly, universal bacterial 16S rRNA genes were PCR amplified using 926F and 1392wR primers and then sequenced on an Illumina MiSeq using 30% PhiX Control v3 (Illumina) and a MiSeq Reagent Kit v3 (600 cycles; Illumina). The following processing steps were then applied to forwards reads using USEARCH (v10.0.240) (Edgar, 2010): 1) primers were removed and the residual sequences were trimmed to 250 bp using fastx_truncate; 2) high-quality sequences were identified using fastq_filter by discarding reads with greater than one expected error (-fastq_maxee=1); 3) duplicate sequences were removed using fastx_uniques; 4) sequences were clustered at 97% similarity into operational taxonomic units (OTU) and potential chimeras were identified and removed using cluster_otus; and 5) an OTU table was generated using otutab with default parameters from the pre-trimmed reads and the OTU representative sequences. SILVA SSU (v128) (Quast et al., 2013) taxonomy was assigned using BLASTN (v2.3.0+) (Zhang et al., 2000) within the feature classifier of QIIME2 (v2017.9) (Boylen et al., 2018). The OTU table was then filtered to remove OTUs classified as chloroplasts, mitochondria, archaea or eukaryotes using the BIOM (McDonald et al., 2012) tool suite. De-novo multiple sequence alignments of the representative OTU sequences were generated using MAFFT (v7.221) (Katoh and Standley, 2013), masked with the alignment mask command of QIIME2 and then used to generate a midpoint-rooted phylogenetic tree using FastTree (v2.1.9) (Price et al., 2010) in QIIME2. OTU tables were then rarefied to 4,850 sequences per sample and alpha diversity metrics were calculated using QIIME2.

### 2.4 Statistical analyses

For statistical analyses, we defined Treatment as the combination of applied substance (none for the control, GO, rGO, aGO, CNTs or graphite) and applied dose (1 ng, 1 µg or 1 mg kg dry soil^−1^). Hence, Treatment was defined as a categorical variable with 16 classes. In order to determine whether the GO, rGO, aGO, CNT and graphite treatments significantly affected the alpha diversity metrics, we used a linear mixed-effects model approach (Pinheiro and Bates, 2004). Treatment (as defined above) and Day, as well as their interaction, were treated as fixed effects, and soil containers (samples) were treated as a random effect to account for the repeated measures. F-tests were applied to assess significance (*P*<0.05), and were implemented in R using the lme4 (Bates et al., 2015) and lmerTest (Kuznetsova, 2017) packages.

Differences in the relative abundances of taxa between samples (beta diversity) were assessed using multivariate generalized linear models using a negative binomial distribution (Warton, 2011). The significance of differences in community composition was determined by comparing the sum-of-likelihood test statistics for the alternative statistical models via a resampling method (Wang et al., 2012) that accounted for the correlation between species and the correlation within the repeated measures taken from the same sample container. These comparisons were implemented in R using the mvabund package (Wang et al., 2012). Taxa whose maximum relative abundance was less than 0.1% were disregarded before statistical analysis. Where an interactive effect of Treatment and Day was significant, post-hoc analyses were undertaken to investigate which Treatments differed on what Days. Where no interactive effect was found, but a main effect of Treatment was, post-hoc analyses focused on which Treatments differed from one another. The Benjamini-Hochberg correction was applied to all post-hoc tests.

## 3. Results

### 3.1 Alpha diversity of soil bacterial communities

Alpha diversity, as represented by the numbers of observed (Sobs) and predicted (Chao1) bacterial taxa, Simpson Diversity Index and Faith’s Phylogenetic Diversity Index (PD), were not significantly influenced by any of the treatments relative to the controls (Fig. S1).

### 3.2 Soil bacterial community composition

Soil bacterial communities were dominated by representatives of the Acidobacteria, Actinobacteria, Armatimonadetes, Bacteroidetes, Candidatus Berkelbacteria, Chlamydiae, Chlorobi, Chloroflexi, Fibrobacteres, Firmicutes, Gemmatimonadetes, Microgenomates, Planctomycetes, Proteobacteria, Saccharibacteria, Spirochaetae and Verrucomicrobia (Fig. S2).

Relative to the no-treatment controls, the composition of bacterial communities was significantly influenced by GO, rGO, aGO, CNTs and Graphite (Table 1; Fig. 1), and these effects differed over time (*P* < 0.003). With three exceptions on Day 14, significant effects were observed for all materials at all doses (Table 1; Table S1). While dose effects were significant, ordination revealed that increasing dose did not lead to consistent directional changes in bacterial community composition (Fig. 1).

**Table 1.** Summary of multivariate GLM post-hoc results (*P* values) computed using mvabund highlighting differences in bacterial community composition between treatments relative to the no-treatment controls within each time point

| Material | Dose | Day |  |  |
| --- | --- | --- | --- | --- |
|  |  | 7 | 14 | 30 |
| Graphite | Low | 0.002** | 0.002** | 0.002** |
|  | Medium | 0.002** | 0.002** | 0.002** |
|  | High | 0.006** | 0.010** | 0.002** |
| GO | Low | 0.002** | 0.207 | 0.003** |
|  | Medium | 0.004** | 0.006** | 0.002** |
|  | High | 0.003** | 0.042* | 0.023* |
| rGO | Low | 0.002** | 0.081 | 0.007** |
|  | Medium | 0.006** | 0.035* | 0.007** |
|  | High | 0.009** | 0.025* | 0.003** |
| aGO | Low | 0.009** | 0.009** | 0.002** |
|  | Medium | 0.003** | 0.034* | 0.006** |
|  | High | 0.005** | 0.053 | 0.006** |
| CNTs | Low | 0.009** | 0.005** | 0.002** |
|  | Medium | 0.004** | 0.024* | 0.002** |
|  | High | 0.003** | 0.003** | 0.010** |

**Fig. 1.**
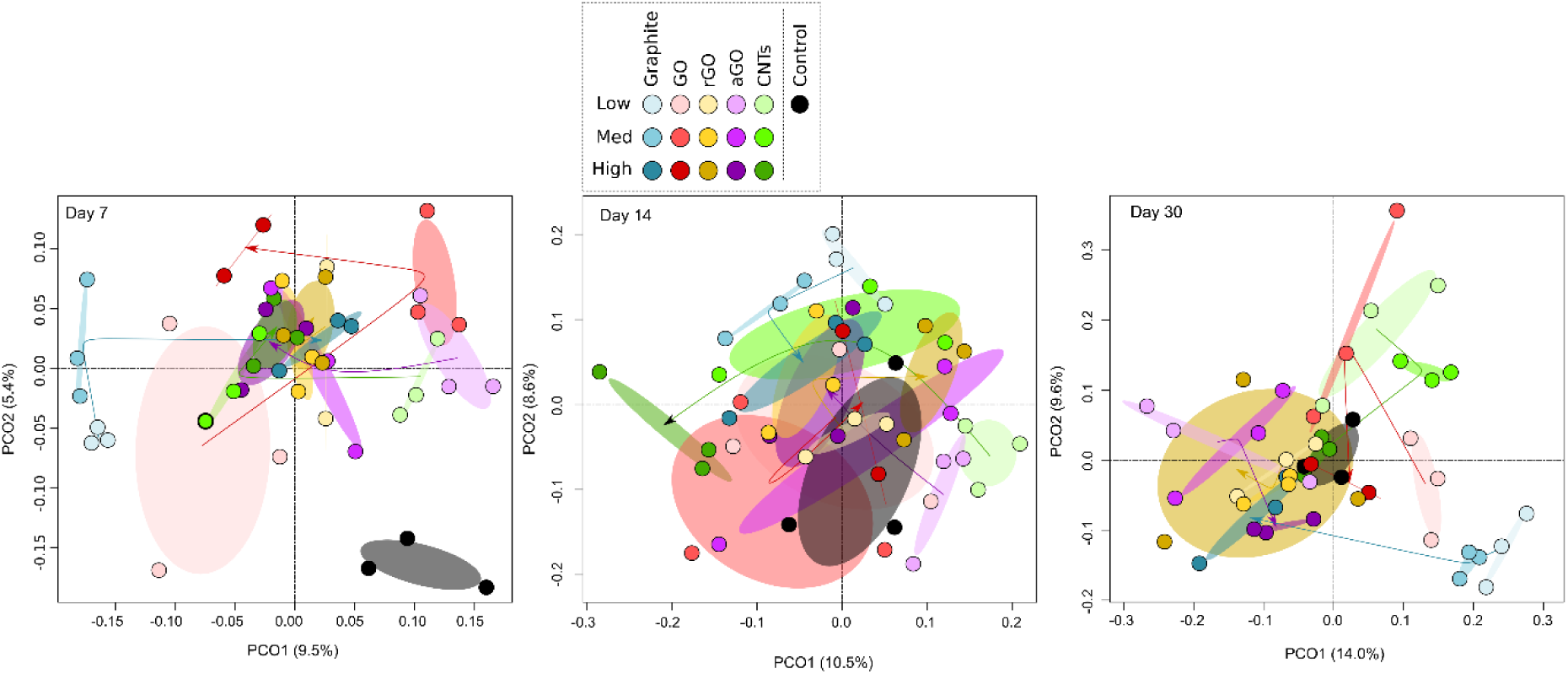
Principal coordinate analysis (PCoA) ordination illustrating differences in the composition of bacterial communities in the no-treatment controls and CNT, GO, rGO, aGO, and graphite amended soils over time. The ellipses represent standard deviations. The arrows for each treatment move from low, through medium, to high dose. They highlight that there were no consistent directions of change in community composition with increasing nano-material or graphite dose. The third replicate for the low dose of rGO on day 7 is outside the plot area (0.22,-0.41).

Relative to Graphite, all nanomaterials led to significant changes in bacterial community composition at all doses, except for aGO at high dose (Table 2). While the effects of most treatments were significant at all time-points, those associated with high concentrations of GO and rGO were significant in later time-points only (Table 2).

**Table 2.**
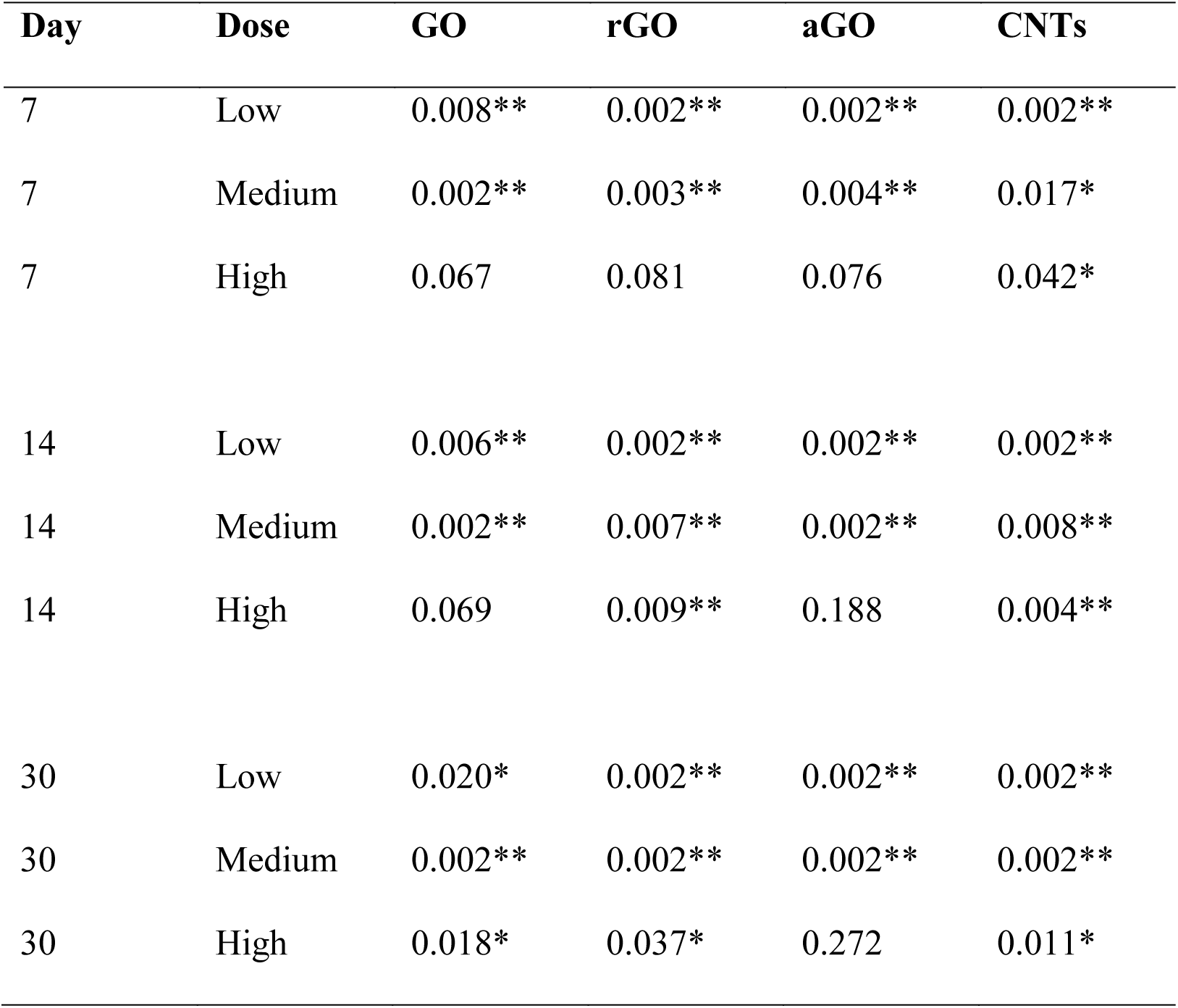
Summary of multivariate GLM post-hoc results (*P* values) computed using mvabund highlighting differences in bacterial community composition between treatments relative to graphite at the same dose within time points

| **Day** | **Dose** | **GO** | **rGO** | **aGO** | **CNTs** |
| --- | --- | --- | --- | --- | --- |
| 7 | Low | 0.008** | 0.002** | 0.002** | 0.002** |
| 7 | Medium | 0.002** | 0.003** | 0.004** | 0.017* |
| 7 | High | 0.067 | 0.081 | 0.076 | 0.042* |
| 14 | Low | 0.006** | 0.002** | 0.002** | 0.002** |
| 14 | Medium | 0.002** | 0.007** | 0.002** | 0.008** |
| 14 | High | 0.069 | 0.009** | 0.188 | 0.004** |
| 30 | Low | 0.020* | 0.002** | 0.002** | 0.002** |
| 30 | Medium | 0.002** | 0.002** | 0.002** | 0.002** |
| 30 | High | 0.018* | 0.037* | 0.272 | 0.011* |

Relative to CNTs, GO and its derivatives led to significant changes in bacterial community composition at all doses, and these effects were apparent in most time-points (Table 3).

**Table 3.** Summary of multivariate GLM post-hoc results (*P* values) computed using mvabund highlighting differences in bacterial community composition between GO and its derivatives (rGO and aGO) relative to CNTs at the same dose

| Day | Dose | GO | rGO | aGO |
| --- | --- | --- | --- | --- |
| 7 | Low | 0.002** | 0.005** | 0.046* |
| 7 | Medium | 0.007** | 0.021* | 0.023* |
| 7 | High | 0.083 | 0.063 | 0.021* |
| 14 | Low | 0.002** | 0.003** | 0.124 |
| 14 | Medium | 0.002** | 0.095 | 0.023* |
| 14 | High | 0.005** | 0.002** | 0.005** |
| 30 | Low | 0.004** | 0.006** | 0.003** |
| 30 | Medium | 0.002** | 0.002** | 0.002** |
| 30 | High | 0.253 | 0.011* | 0.006** |

Lastly, relative to GO, the rGO and aGO treatments led to significant changes in bacterial community composition at all doses (Table 4). These effects were apparent for both materials at all doses (except high rGO) at the beginning and end of the experiment, but were variable on Day 14 (Table 4).

**Table 4.** Summary of multivariate GLM post-hoc results (*P* values) computed using mvabund highlighting differences in bacterial community composition between GO and its derivatives (rGO and aGO) at the same dose

| **Day** | **Dose** | **rGO** | **aGO** |
| --- | --- | --- | --- |
| 7 | Low | 0.002** | 0.003** |
| 7 | Medium | 0.009** | 0.010** |
| 7 | High | 0.055 | 0.037* |
| 14 | Low | 0.104 | 0.003** |
| 14 | Medium | 0.011* | 0.009** |
| 14 | High | 0.080 | 0.108 |
| 30 | Low | 0.003** | 0.002** |
| 30 | Medium | 0.002** | 0.003** |
| 30 | High | 0.022* | 0.012* |

The 100 OTUs that were most strongly associated with differences in community composition between treatments were obtained from the multivariate GLMs and assessed independently using univariate GLM models. Of these, 39 were found to differ significantly from the no-treatment controls in at least one treatment combination after Benjamini-Hochberg correction for multiple comparisons (Fig. 2).

**Fig. 2.**
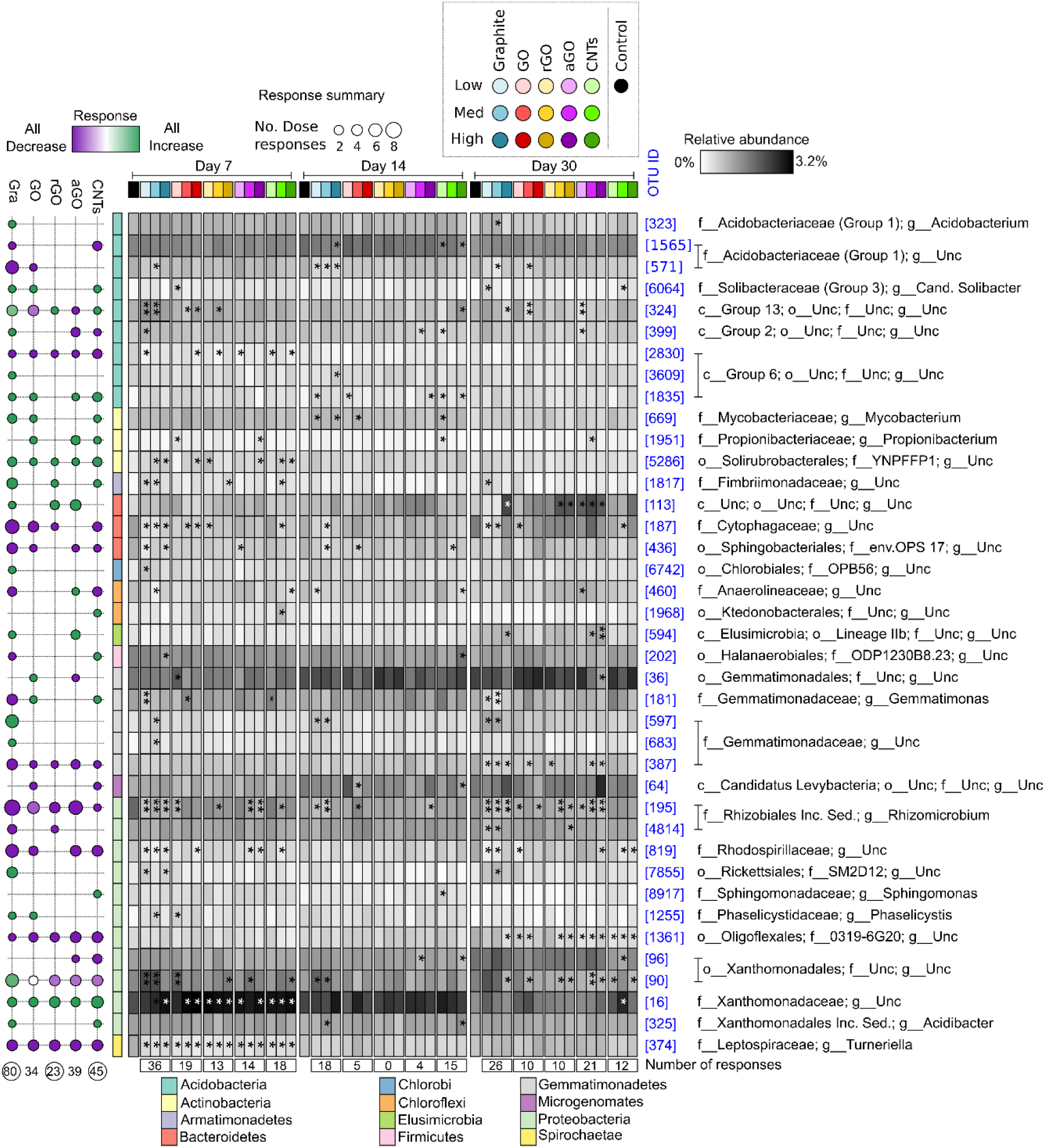
Heatmap of the relative abundances of 39 bacterial OTUs that differed significantly from the control in at least one treatment combination. The asterisks highlight which treatments differ significantly from the no-treatment controls on each day (*P* < 0.05*, *P* < 0.01**, *P* < 0.001***). Each column of the heatmap represents the mean relative abundance of each treatment (n = 3). The bubble-plot on the left summarizes the number (circle size) of nanocarbon or graphite treatments that an OTU responded to relative to the no-treatment controls, and of these how many manifested as increases or decreases in relative abundance (circle color). The numbers below the bubble plot and heatmap show the total numbers of significant responses to a particular treatment relative to the control. The OTU IDs are consistent throughout the manuscript. The phylum of each OTU is indicated by the colors on the left of the heatmap and the affiliations associated with each color are shown at the bottom.

Six OTUs responded exclusively to graphite, an *Acidobacterium* (OTU 323; Acidobacteria), two members of the *Gemmatimonadaceae* (OTU 597, OTU 683; Gemmatimonadetes), a Rickettsiales (OTU 7855, Proteobacteria), Acidobacteria of subgroup 6 (OTU 3609) and a member of the Chlorobiales (OTU 6742, Chlorobi) (Fig. 2). Two OTUs responded exclusively to CNTs; these were a member of the Ktedonobacterales (OTU 1968, Chloroflexi) and a member of the *Sphingomonas* (OTU 8917, Proteobacteria). All of these OTUs responded with increases in relative abundance when exposed to graphite or CNTs (Fig. 2). A further four OTUs responded to some combination of multiple nanocarbon materials but did not respond to graphite addition. These included a member of the *Propionibacteriaceae* (OTU 1951, Actinobacteria) which increased in relative abundance; a Levybacteria (OTU 64, Microgenomates) and a Xanthomonadales (OTU 96, Proteobacteria) population, which decreased in relative abundance; and a member of the Gemmatimonadales (OTU 36, Gemmatimonadetes), which increased in response to GO and decreased in response to aGO. A further nine OTUs responded to all nanomaterials, while a further 18 responded to the addition of graphite, as well as at least one nanomaterial (Fig. 2).

## 4. Discussion

Previously, we reported that graphite and GO influence the composition of soil microbial communities at concentrations deemed realistic by models of release rates for CNTs and other nanomaterials into soils (Forstner et al., 2019). Here, we extend these findings by demonstrating that relative to no-treatment controls, graphite and GO, the composition of bacterial communities is also significantly influenced by the addition of CNTs, rGO and aGO. Irrespective of concentration, all nanomaterials led to shifts in community composition that were of similar magnitude to those associated with graphite, albeit with differences in the taxa affected. Despite the significant changes to bacterial community composition the alpha diversity of bacterial communities remained unaffected by all treatments (Fig. S1).

Previous studies examining the effects of nanocarbon materials have demonstrated a wide variety of effects on bacterial communities both in culture and in soils. Most of these studies have been focused on either CNTs or GO and have demonstrated an array of effects ranging from inhibition of enzyme activities (Chung et al., 2011, 2015; Jin et al., 2013; Xiong et al., 2018) to reductions of biomass (Chung et al., 2011; Jin et al., 2013, 2014) and shifts in microbial community composition (Du et al., 2015; Forstner et al., 2019; Ge et al., 2016; Jin et al., 2014; Xiong et al., 2018). Availability of data regarding the effects of rGO and aGO on soil microbes is minimal, however. rGO has been shown to have some antibacterial properties in pure culture (Guo et al., 2017; Gurunathan et al., 2012; Liu et al., 2011), whereas no research exists on the effects of aGO on microbes. While some of the studies on CNTs and GO have examined soil bacterial communities (Chung et al., 2011, 2015; Du et al., 2015; Ge et al., 2016; Jin et al., 2013, 2014; Khodakovskaya et al., 2013; Rodrigues et al., 2013; Shrestha et al., 2013; Xiong et al., 2018), these have used doses that are ≥6,250 times the estimated annual rate of accumulation for CNTs (Sun et al., 2014). In our previous work, we observed significant and diverse shifts in soil microbial community composition after application of GO at realistic exposure concentrations (Forstner et al., 2019), highlighting that the effects of nanocarbon on soil microbial communities may persist in some manner even at realistic exposure rates. However, while this work demonstrated similarities between the effects of GO and graphite, it did not compare these effects against those elicited by other carbon nanomaterials.

Our study illustrates that the degree of bacterial community compositional changes did not increase with the application rate of nanocarbon materials but varied over time and we observed fewer significant differences between treatments at high dose than at medium or low dose (Table 2, 3, 4, Fig. 1). This suggests that even low rates of nanocarbon application in the range of ng-mg kg^−1^ can induce significant compositional changes in soil bacterial communities and these changes can persist for at least 30 days and that at mg kg^−1^ dose the changes elicited by different nanocarbon materials converge somewhat. We propose that the absence of a ‘linear’ dose response to nanocarbon addition is, at least partially, due to inhibition of the free movement of particles and the availability of sharp edges as well as clean nanocarbon surfaces. Past studies have identified mechanisms which can reduce the antibacterial effects of nanocarbon materials; agglomeration (Dreyer et al., 2010; Liu et al., 2009), covering of nanocarbon structures with contaminants (Hui et al., 2014), as well as interactions with components of the soil matrix (Chen et al., 2018; Jaisi and Elimelech, 2009; Lu et al., 2017). For example, rGO (Sengupta et al., 2019) and CNTs (Hartono et al., 2018) can damage bacterial cell membranes through mechanical stress and piercing (Hartono et al., 2018; Sengupta et al., 2019). However, this interaction is dependent on both the mobility of particles as well as the availability of clean edges on nano-sheets and tubes.

While all amendments elicited significant shifts in soil bacterial community composition, a minority of affected taxa responded exclusively to one material. Importantly, 69% of discriminating taxa responded both to graphite and at least one nanomaterial, with a third of these responding to all materials. Furthermore, 15% of OTUs responded only to graphite, thereby supplying the largest group of exclusively responding OTUs. The only other group of taxa responding in such an exclusive manner were the 5% of taxa responding only to CNTs. None of the taxa were solely affected by GO and its derivatives during the experiment. A further 10% of discriminating taxa responded to multiple nanocarbon materials but not to graphite.

In summary, most populations responded to graphite, with many of these also responding to one or more nanomaterials and 75% of exclusively responding taxa did so in response to graphite addition. The only nanomaterial that elicited responses not seen in other materials were the CNTs. This is most likely a result of their tubular nature, potentially resulting in interactions with bacteria that are in some manner, such as by shape or structure, shielded against interactions with nano-sheets. Such shielding effects have been observed for GO and rGO in pure cultures (Sengupta et al., 2019). The number of relative increases and decreases in abundance were approximately equal across materials. This is consistent with previous research which has reported variable effects of GO (Akhavan and Ghaderi, 2010; Chung et al., 2015; Du et al., 2015; Forstner et al., 2019; Liu et al., 2011; Ruiz et al., 2011; Xiong et al., 2018) and CNTs (Chung et al., 2011; Ge et al., 2016; Hartono et al., 2018; Jin et al., 2014, 2013, Kang et al., 2009, 2008; Khodakovskaya et al., 2013; Rodrigues et al., 2013; Shrestha et al., 2013) on bacterial isolates as well as soil bacterial communities and has reported that CNTs may cause changes similar in magnitude to reference materials such as biochar and carbon black (Ge et al., 2016) albeit at much greater doses than can be deemed realistic (Ge et al., 2016; Sun et al., 2014).

Previous work has highlighted the effects of very high (10 g kg^−1^) CNT applications on various taxa (Shrestha et al., 2013). Shrestha *et al.* (2013) highlighted increases in the relative abundance of, among others, *Rhodococcus* and *Cellulomonas* as well as decreases in *Holophaga* and *Derxia* (Shrestha et al., 2013). We did not observe any changes in any of the taxa identified by Shrestha et al. (2013) under the addition of any of the treatments. This is likely due to the up to a 10-billion-fold difference in doses applied between the studies. Our examination of OTUs responding to treatments in univariate GLMs revealed that the number of responses elicited by nanomaterials was exceeded by responses to graphite in all time-points with rGO addition consistently resulting in the smallest number of responses; with not a single taxon responding at 14 days. The number of responses elicited by CNTs decreased over time, while aGO experienced a drastic increase from 4 to 21 responses between day 14 and 30. This resulted in aGO being the only material that elicited more responses at the end than at the beginning of the experiment (Fig. 2). Further research is required in order to determine if the mobilization of ammonia bound to GO sheets is the source of this discrepancy.

## 5. Conclusions

With increasing utilization of nanocarbon materials, it is becoming ever more pressing to examine their ultimate environmental effects. Our study demonstrates that CNTs, rGO and aGO can significantly influence bacterial community composition even at doses as low as one ng kg^−1^ and that these effects significantly differ from not only a no-treatment control but also from those elicited by graphite and unmodified GO sheets. This highlights not only the importance of examining nano-carbon materials as a whole; but also reveals a need to examine each new material and derivative of existing materials in order to determine their environmental impacts on the soil microbial communities that underpin essential ecosystem services.

## Data availability

All sequences have been deposited to the Sequence Read Archive (SRA) under BioProject accession number PRJNA515098

## Acknowledgements

The authors gratefully acknowledge financial support from The University of Queensland for an Early Career Researcher Award to PGD. CF gratefully acknowledges funding from the Australian Government’s Department of Education and Training in the form of an Australian Government Research Training Program Scholarship administered by The University of Queensland. PMK is the recipient of an Australian Research Council (ARC) Future Fellowship (ARC FT120100277).

