## Supplementary information for "Effects of carbon nanotubes and derivatives of graphene oxide on soil bacterial diversity"

**Fig. S1** Barcharts of observed alpha diversity metrics by treatment..........................................2

**Fig. S2** Relative abundance of bacterial phyla...........................................................................3

**Table S1** Pairwise comparisons between doses for bacteria.....................................................4

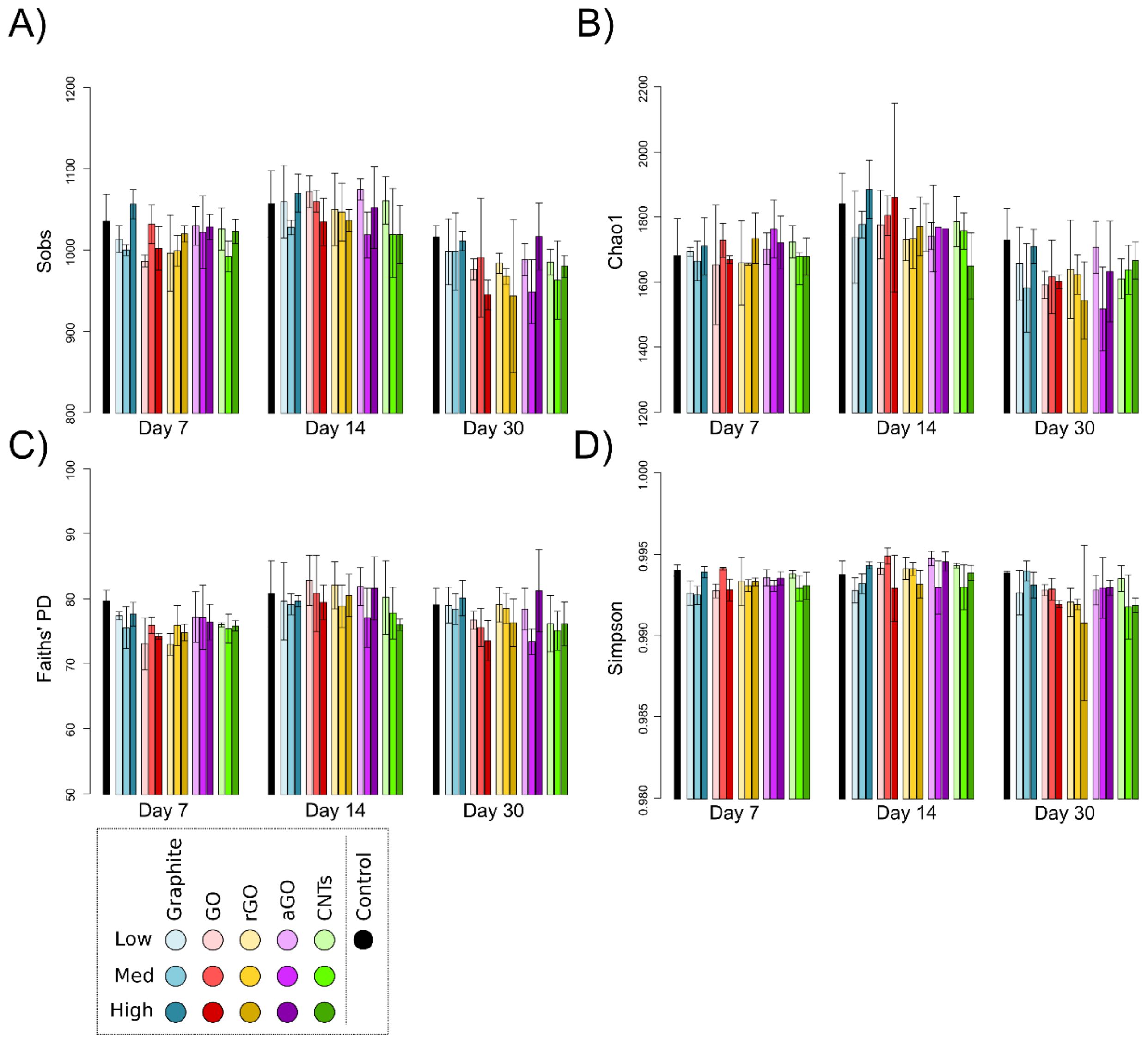


**Fig. S1** The alpha diversity of bacterial communities over time: A) the numbers of observed (Sobs) bacterial OTUs; B) the numbers of predicted (Chao1) bacterial OTUs; C) Faith's Phylogenetic Diversity Index; and D) Simpson's Diversity Index. Error bars represent standard deviations. None of the treatments differed significantly from the controls.

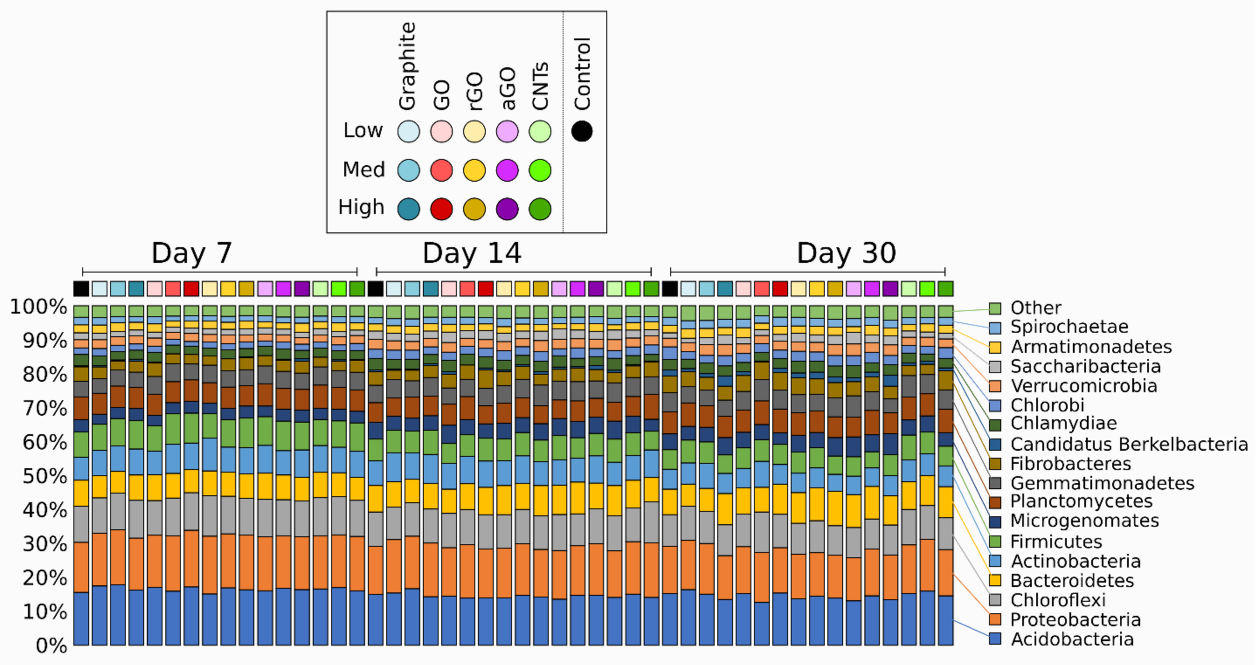


**Fig. S2** The relative abundances of bacteria phyla by treatment over time. All phyla representing <1% relative abundance are combined as "Other".

**Table S1** Summary of multivariate GLM post-hoc results (*P* values) computed using mvabund highlighting differences in bacterial community composition between different doses of the same material over time

| **Material** | **Dose 1** | **Dose 2** | **Day** | | |
|---|---|---|---|---|---|
| | | | 7 | 14 | 30 |
| Graphite | Low | Medium | 0.042* | 0.022* | 0.107 |
| | Low | High | 0.005** | 0.007** | 0.002** |
| | Medium | High | 0.002** | 0.002** | 0.002** |
| GO | Low | Medium | 0.002** | 0.006** | 0.002** |
| | Low | High | 0.020* | 0.107 | 0.069 |
| | Medium | High | 0.017* | 0.009** | 0.008** |
| rGO | Low | Medium | 0.016* | 0.131 | 0.235 |
| | Low | High | 0.014* | 0.047* | 0.058 |
| | Medium | High | 0.074 | 0.021* | 0.058 |
| aGO | Low | Medium | 0.024* | 0.021* | 0.006** |
| | Low | High | 0.011* | 0.006** | 0.004** |
| | Medium | High | 0.036* | 0.037* | 0.009** |
| CNTs | Low | Medium | 0.006** | 0.002** | 0.030* |
| | Low | High | 0.018* | 0.002** | 0.005** |
| | Medium | High | 0.048* | 0.002** | 0.003** |
